# Predicting future conservation areas while avoiding competition in two alpine urodele amphibians severely threatened by climate change

**DOI:** 10.1101/2023.03.24.534075

**Authors:** Nicolas Dubos, Antoine Havard, Angelica Crottini, Daniele Seglie, Franco Andreone

## Abstract

Climate change will cause important declines in species distributions, especially when living at high altitudes. The Critically Endangered *Salamandra lanzai* from SW Alps may be severely exposed to future climate change effects and its suitable climate may shrink or shift. Another Alpine salamander (*S. atra*) is present in the region, which in case of spatial overlap may represent a competitor for *S. lanzai*. It is urgent to estimate the effect of future climate change on these species and identify priority areas for conservation while accounting for competition between both species. With a Species Distribution Modelling (SDM) approach, we projected the current and future climate suitability of both salamander species. We accounted for uncertainty related to the methods (model replicates) and climate projections (data source, global circulation model and scenario) to provide a consensus map for practitioners. This map also takes into account potential competition with *S. atra* by penalizing the suitability scores of S. *lanzai* by the scores of *S. atra*. We predict a severe effect of climate change on both species. Most of the current habitats are projected to become largely unsuitable by 2070, regardless of the climatology and scenario. We identified important spatial disagreements between projections based on different data sources, mostly due to precipitation projections and daily temperature variation. This highlights the need to account for multiple climatologies in mountainous environments. Both species’ habitats are highly fragmented, which is expected to prevent distributional shifts through natural dispersion. We suggest to explore the possibility of translocation for the most threatened populations and simultaneously develop captive breeding programs. Biotic interactions are rarely accounted for in SDMs, and we encourage the documentation of species with similar ecological requirements to improve the relevance of SDMs for future conservation planning.

## Introduction

Climate change is one of the main threats to montane species, especially amphibians (Raxworthy et al., 2008; Mccain & Colwell, 2011). Warming temperatures will likely cause upslope shifts in species distributions, potentially leading to extinction (Freeman & Class Freeman, 2014; Rana et al., 2021). The effect of climate change varies along latitude (Tewksbury et al. 2008; Dubos et al. 2019), which particularly applies to montane species, with temperate species less likely to respond negatively than tropical ones (Freeman et al. 2021). Upslope shifts may not always reduce the amount of suitable area in some systems but also bring newly available regions, thus increasing the availability of suitable habitats (Elsen & Tingley, 2015). Hence there is a need to compile climate change predictions for mountainous environments and quantify the amount of suitable environments in the future.

Species distribution models (SDMs) can be used to identify the most suitable conditions for a given species, and forecast potential future geographical shifts in these conditions. SDMs are commonly used for testing ecological hypotheses (Anderson et al., 2009; Dubos, Augros, et al., 2022), assessing alien species invasion risks (Bellard et al., 2013; Dubos, Fieldsend, et al., 2022; Lanner et al., 2022), forecasting the potential effect of environmental change (Araújo et al., 2005), and supporting conservation and management efforts (Schwartz, 2012; Leroy et al., 2014; Dubos, Lenormand, et al., 2022). In conservation application, SDMs can provide maps of projected future environmental suitability at the resolution of environmental data. These maps can be used as a tool for practitioners to seek suitable habitats or microhabitats within the identified suitable pixels and propose priority areas for conservation actions (e.g. for the identification of potential translocation sites, or for protection status).

Biotic interactions are rarely accounted for in SDMs because they are highly demanding in term of data and analysis. For a given species, suitable climatic conditions may shift with climate change, and these conditions may overlap with that of another species sharing similar ecological requirements. The sites identified as suitable in the future may represent candidate sites for conservation action for both species, but would expose them to competition. Hence, there is a need to identify the best candidate sites for either species while mutually excluding them.

We studied the effect of climate change on the potential distribution of two alpine salamanders with similar ecological adaptations, Lanza’s salamander (*Salamandra lanzai*) and alpine salamander (*S. atra*). Both these species are typical of montane environments and evolved an aplacental viviparity (with full embryonic development occurring within their body), that allowed them to complete their reproductive cycle independently from the occurrence of water bodies. This is likely an adaptation to cold climate at high temperatures (Ma et al., 2018). In both species, several populations are highly isolated due to both their past biogeographic histories and more recent anthropogenic landscape modification (Tessa, Crottini, & Andreone, 2007). Current patchy distribution suggests that individuals from these isolated populations will likely not be able to follow the shifting climatic conditions through dispersal (Andreone et al. 2004). In this study we identified the climatic predictors explaining the current distribution of *S. lanzai* and *S. atra* using an SDM approach. We expect a strong effect of climate change on both species, especially for *S. lanzai* due to its narrow distribution (Platts et al., 2014; Dubos, et al., 2022). Due to its conservation status (Critically Endangered; IUCN SSC Amphibian Specialist Group, 2022), we focus on *S. lanzai* for conservation considerations. We projected both species climate suitability on current and future (2070) conditions in the Alps in order to (1) quantify the effect of climate change on both species’ distributions and (2) identify priority areas for the conservation of *S. lanzai* while accounting for competition.

## Methods

### Occurrence data

We obtained 1991 and 841 unique locality records for *S. lanzai* and *S. atra*, respectively. These have been retrieved from iNaturalist (https://www.inaturalist.org/) and Parc Naturel Régional du Queyras (https://www.pnr-queyras.fr/). To avoid pseudo-replication, we resampled one locality record per pixel of climate data (i.e. data thinning; Steen et al. 2021). At the resolution of 1 × 1 km, this led to a final sample of n = 104 or n = 102 presence points for *S. lanzai* after resampling on CHELSA data, or with Worldclim, respectively. The final sample for *S. atra* was n = 711 presence points based on CHELSA (710 with Worldclim).

### Climate data

We used six uncorrelated bioclimatic variables (Pearson’s r < 0.7; Dormann et al. 2013) for which we had hypotheses of causal relationship with both species’ biology (Fourcade, Besnard, & Secondi, 2018; Dubos, Préau, et al., 2022) at 30 arc seconds resolution (approximately 1 km) for the current and future (2070) climate from two sources: CHELSA (Karger et al., 2017) and Worldclim global climate data (Fick & Hijmans, 2017). These data sources used different methods to compute the climatologies. Worldclim is based on interpolated data with elevation and distance to the coast as predictors in addition to satellite data, while CHELSA is based on statistical downscaling for temperature, and precipitation estimations incorporating orographic factors (i.e., wind fields, valley exposition, boundary layer height). We describe the causality of the relationship between predictors and species’ ecology in Table 1. The six variables retained were annual mean temperature (Bio1, which was collinear with summer and winter mean temperatures), autumn mean temperature (Bio8), isothermality (Bio3), temperature seasonality (Bio4), precipitation of the driest month (Bio14), and summer precipitation (Bio18). We used a statistical process to select the most relevant predictors (elbow criterion for the relative importance, as measured by the randomForest R package; Breiman 2001). We did not include precipitation seasonality (Bio15) despite this variable is often selected in mountainous areas (e.g. De Marchi et al. 2020), because its causal relationship with the biology of alpine salamanders is not clear. To consider the uncertainty related to future climate projections, we used three Global Circulation Models (GCMs; i.e., BCC-CSM1-1, MIROC5, and HadGEM2-AO) and two greenhouse gas emission scenarios (the most optimistic RCP26 and the most pessimistic RCP85).

**Table 1.** Selected variables and ecological hypotheses of causality for the presence of *Salamandra lanzai* and *S. atra* in the Alps. We give the relative importance of each variable as measured by the Random Forest algorithm (mean decrease in Gini index). Predictors with the highest relative importance were included in the models (in bold for each species, selected on the basis of elbow criterion).

| Bioclimatic variable | Relative importance |  | Ecological interpretation | Reference |
| --- | --- | --- | --- | --- |
|  | *S. lanzai* | *S. atra* |  |  |
| Bio1 (annual mean T) | **Worldclim: 0.24**<br>**CHELSA: 0.23** | **Worldclim: 138**<br>**CHELSA: 134** | Summer activity | Ribéron and Miaud 1999; Kissel et al. 2019 |
| Bio3 (isothermality) | **Worldclim: 0.15**<br>**CHELSA: 0.16** | Worldclim: 105;<br>CHELSA: 78 | Higher metabolic cost in very cold nights | Authors' hypothesis |
| Bio4 (T seasonality) | **Worldclim: 0.07**<br>**CHELSA: 0.12** | **Worldclim: 124**<br>**CHELSA: 134** | Higher metabolic cost in highly variable years, driven by hot winters | Kissel et al. 2019 |
| Bio18 (P of warmest quarter) | Worldclim: 0.12<br>CHELSA: 0.05 | **Worldclim: 106**<br>**CHELSA: 117** | Summer activity, food availability, larval dessication risk | Ribéron and Miaud 1999; Kissel et al. 2019; (Romano et al. 2021) |
| Bio19 (P of coldest quarter) | **Worldclim: 0.24**<br>**CHELSA: 0.26** | **Worldclim: 96**<br>**CHELSA: 108** | Overall habitat moisture | Miaud et al. 2001; Andreone et al. 2004; Romano et al. 2021 |

### Land-use data

We accounted for habitat requirements by filtering all unsuitable habitats from our model projections. We used a filter instead of integrating habitat predictors to our models because (1) the species are strongly dependent on given habitat types (hence, removing unsuitable habitats is realistic), (2) this improves statistical parsimony and (3) final projections benefit from finer resolution (climate projections were resampled to fit land-use data). We obtained high-resolution land cover categories from Corine Land Cover 2018 (100 m resolution) derived from remote sensing (Copernicus Service Information, 2022). We assume agricultural and urbanized areas to be unsuitable to the presence of salamanders (Andreone et al., 2004; Geiger, 2006; Reinthaler-Lottermoser et al., 2010; Romano et al., 2018). These habitat types were removed from final projections, only to keep habitats that are used by both species, i.e. coniferous forests, natural open areas and bare rock (Andreone, De Michelis, & Clima, 1999; Ribéron & Miaud, 1999; Miaud et al., 2001; Bonato & Fracasso, 2003, 2014; Andreone et al., 2004).

### Ecological niche modelling

We modelled and projected climatic niches using highly performing single tuned algorithm (Valavi et al. 2021), i.e. Random Forest down-sampled (hereafter RF down-sampled, i.e. RF parametrised to deal with a large number of background samples and few presence records; Prasad et al. 2006). We set RF down-sampled to run for 1000 bootstrap samples/trees, with 5 cross validation runs and 5 sets of 10,000 pseudo-absences. Since data were strongly spatially aggregated, we did not use a spatial partitioning for cross validation runs (Valavi et al. 2019), but a random partitioning instead. We applied a sample bias correction technique consisting in a selection of pseudo-absence points that imitates the spatial bias of presence points (Phillips et al. 2009). To do so, we used a null geographic model, i.e. a map of the density of unthinned presence points, which we use as a probability weight for pseudo-absence selection.

We determined the background extent following a hypothesis of past dispersal, accounting for the Equilibrium hypothesis, and minimizing the surface to better account for sample bias (Vollering et al., 2019). We assume both alpine salamanders were able to disperse in the whole Alps in the past, hence both backgrounds encompassed the Alps and their surroundings. To account for Equilibrium, we mutually excluded the regions where the putative competitor is present, i.e. the Central and Eastern Alps for *S. lanzai* and South-Western Alps for *S. atra*. The fire salamander *Salamandra salamandra* is present at the foot of the Alps. However, we did not exclude the non-mountainous areas at the foot of the Alps because *S. atra* and *S. salamandra* appear not to be in competition, or competition does not lead to spatial segregation (Werner, Lötters, & Schmidt, 2014).

We evaluated model performance using the Boyce Index (Hirzel et al., 2006). An index of 1 suggests that suitability models were perfectly able to predict presence points, a null value suggests that models are no better than random and a negative value implies a counter prediction. All models for which Boyce index < 0.5 were discarded. We present projections of the mean suitability obtained across all highly performing models.

### Conservation application: Accounting for uncertainty and competition

To provide sound conservation guidelines, we built a consensus map of the mean predictions across all highly performing models, discounting the standard deviation (mean – SD; Kujala et al. 2013; Dubos et al. 2022a). Final ‘consensus’ maps represent the areas that were identified as suitable with the lowest disagreement between model replicates, hence the best candidate sites for conservation action.

Priority areas for the conservation of *S. lanzai* may fall into areas occupied by *S. atra* in Central and Eastern Alps, which may expose both species to competition and reduce success probability for possible conservation efforts (e.g. translocation). To avoid potential sympatry and hence, competition between *S. lanzai* and *S. atra*, we excluded areas that are suitable to *S. atra* from the consensus map of *S. lanzai*.

Similarly, we subtracted the consensus suitability of *S. atra* to the consensus suitability of *S. lanzai*, with the obtained result to be interpreted as a priority map for conservation efforts for *S. lanzai* in Central and Eastern Alps.

## Results

### Ecological niche models

We selected four climatic predictors for either species. For *S. lanzai*, we retained bio1, bio3, bio4 and bio19. For *S. atra*, we kept bio1, bio4, bio18 and bio19 (Fig. S1—S4). For *S. lanzai*, models performed well (i.e. suitability scores are high at presence points), with an average Boyce index of 0.67 (CHELSA; 5 models discarded) and 0.84 (Worldclim; no model discarded) for each climate data, respectively. For *S. atra*, model performance was very high (CHELSA: mean Boyce = 0.98; Worldclim: mean Boyce = 0.99; no model discarded).

For *S. lanzai*, current predictions identified different areas with suitable conditions depending on the source of climate data. With Worldclim, suitable conditions can be found further south of its current distribution (Fig. 1). On the contrary with CHELSA, suitable conditions extend to the north of its current distribution (Fig. 2). For *S. atra*, suitable conditions are consistently found across the northern and eastern part of the Alps with both climate data sources (Fig. 3, 4).

**Fig. 1.**
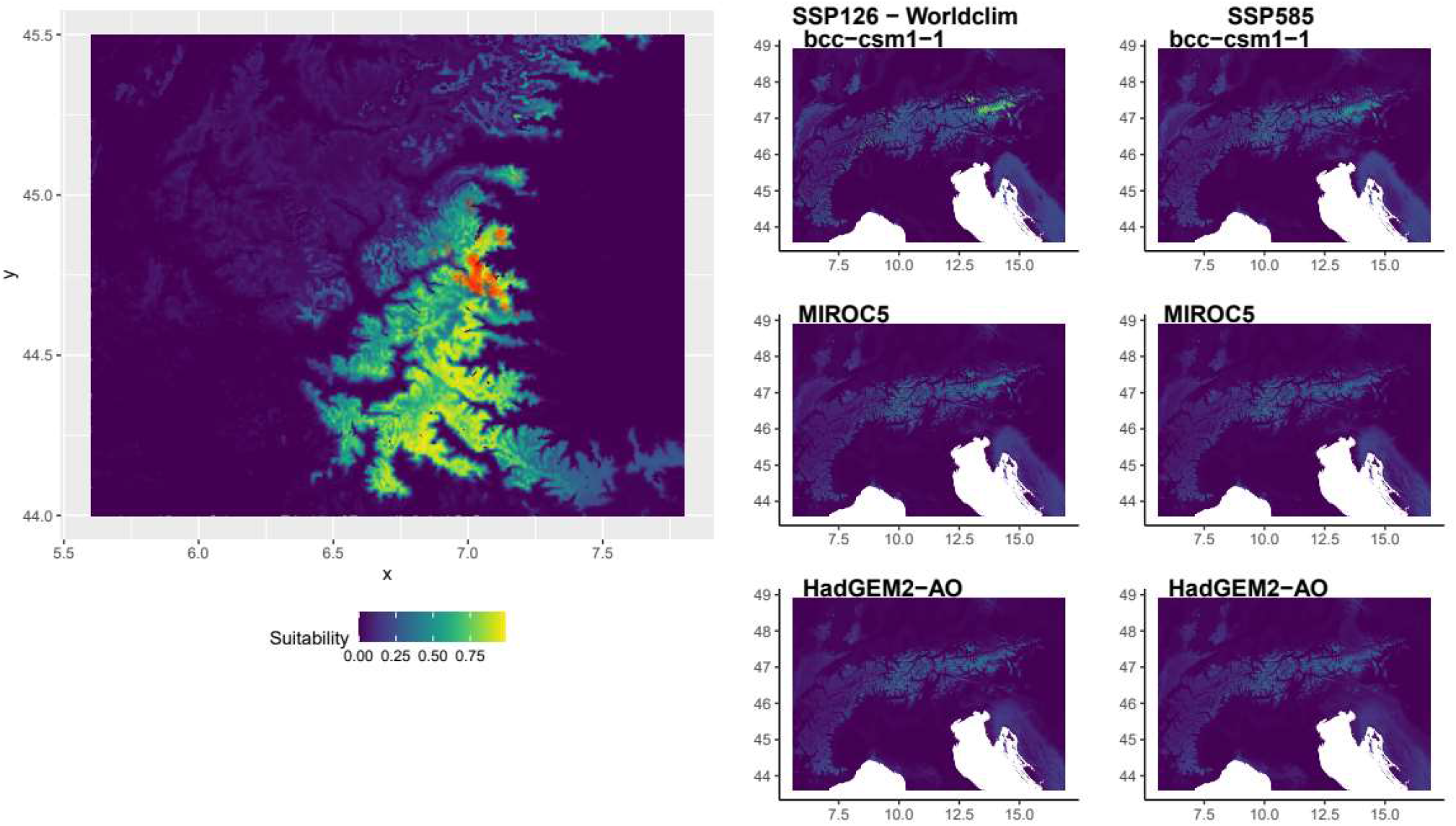
Environmental suitability for *Salamandra lanzai* based on Worldclim climatologies. Left: current; Right: future (2070). Future projections include two emission scenarios (SSP126 and SSP585) and three GCMs.

**Fig. 2.**
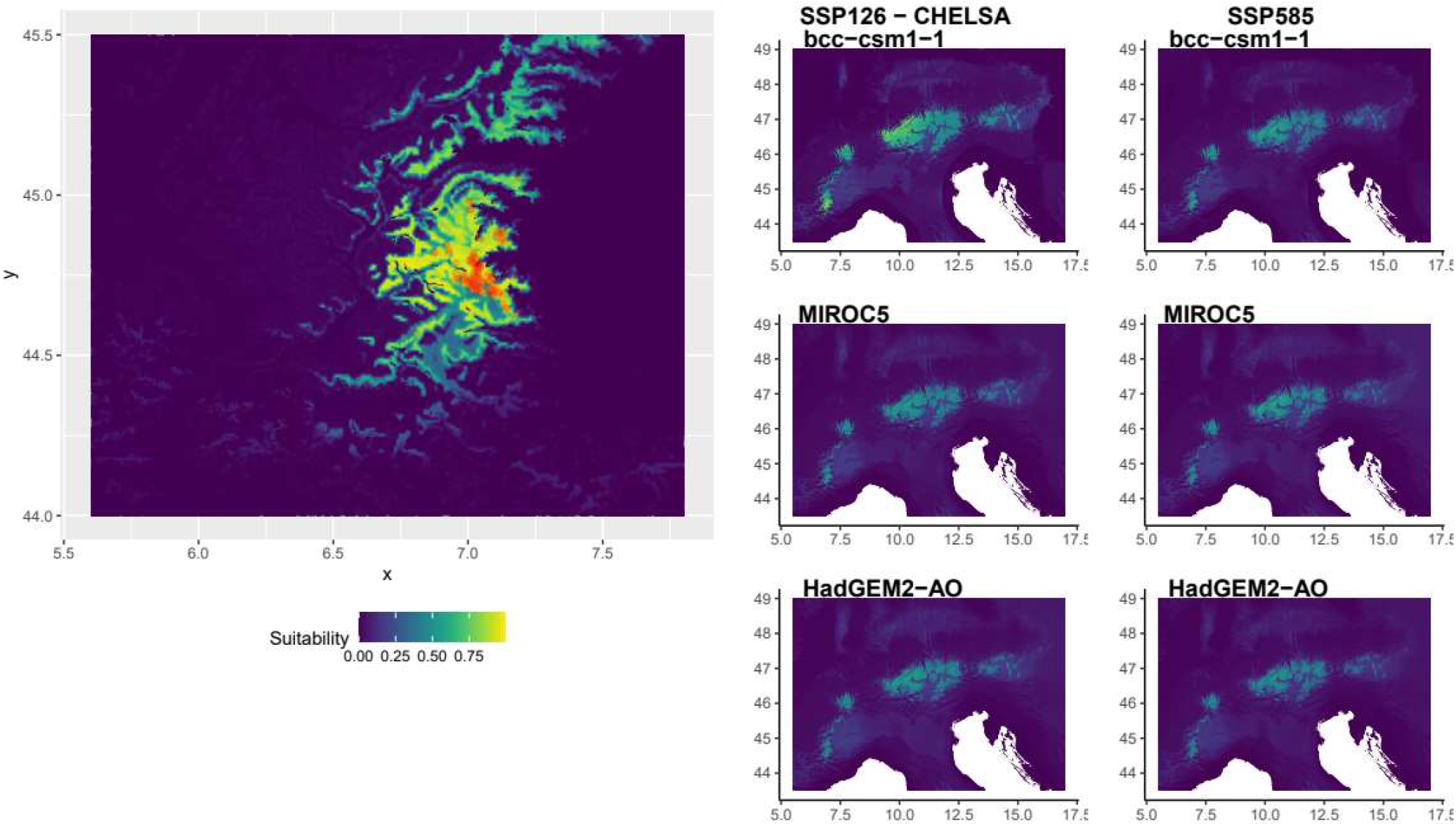
Environmental suitability for *Salamandra lanzai* based on CHELSA climatologies. Left, current; Right, future (2070). Future projections include two emission scenarios (SSP126 and SSP585) and three GCMs.

**Fig. 3.**
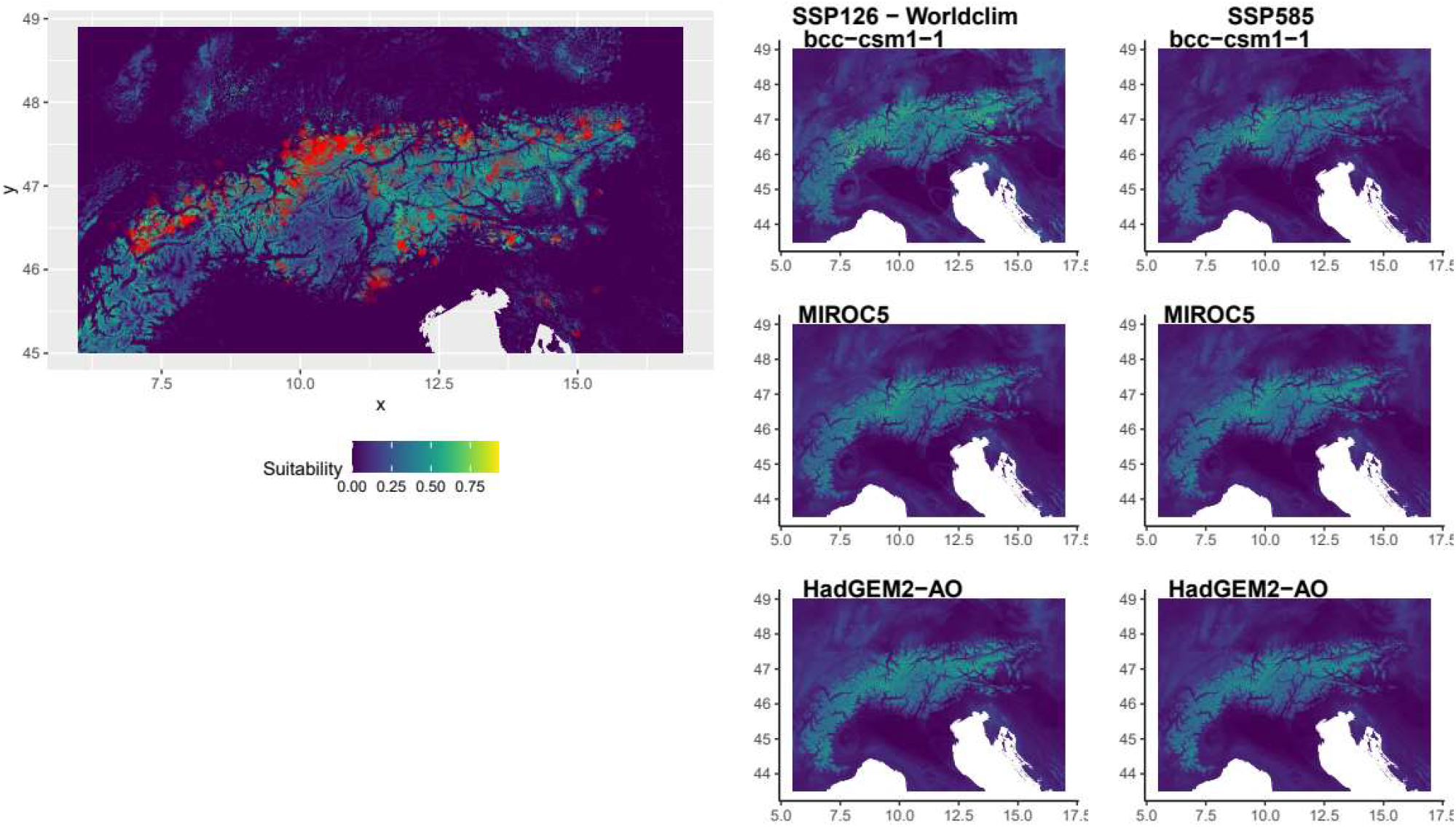
Environmental suitability for *Salamandra atra* based on Worldclim climatologies. Left, current; Right, future (2070). Future projections include two emission scenarios (SSP126 and SSP585) and three GCMs.

**Fig. 4.**
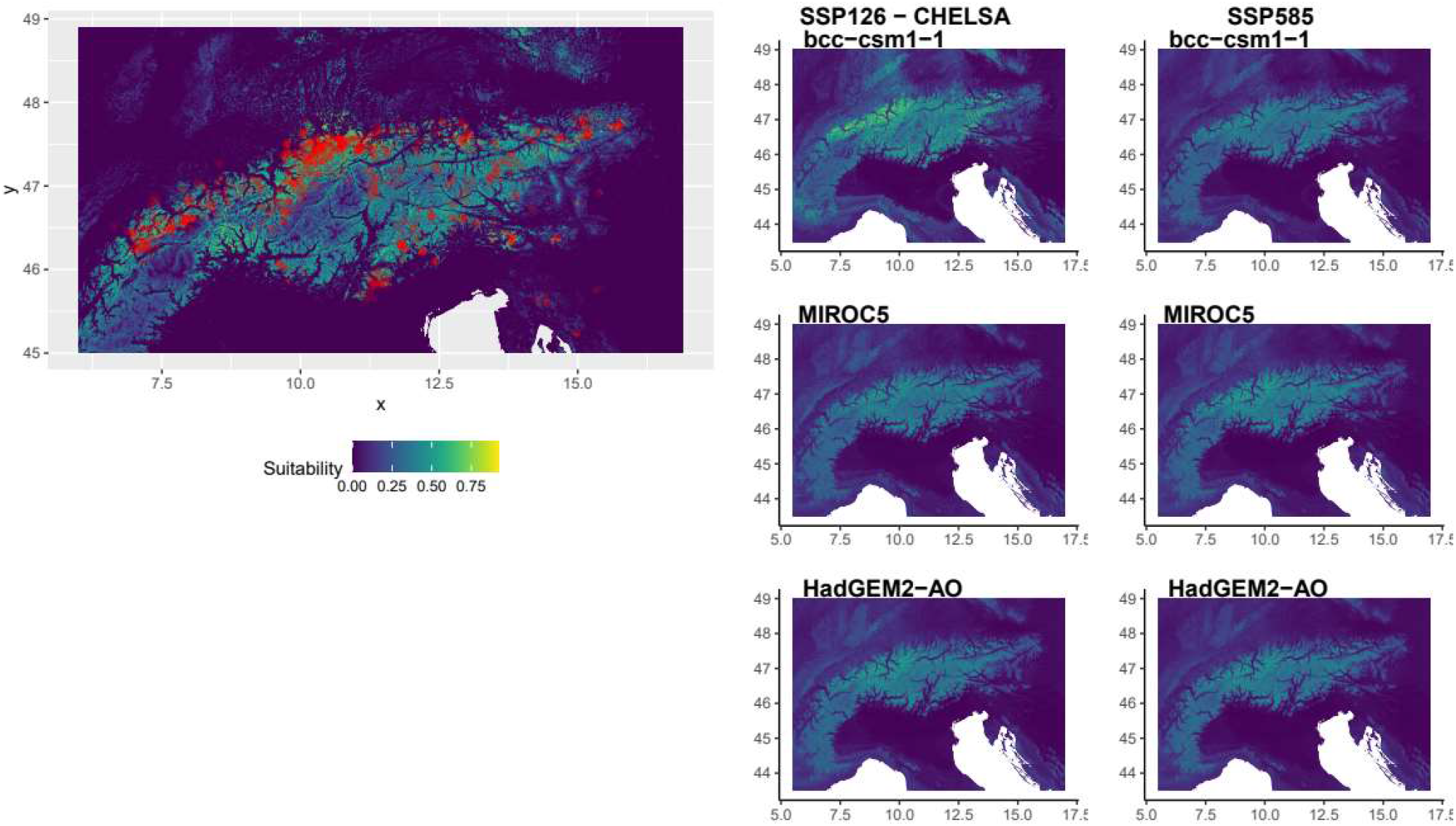
Environmental suitability for *Salamandra atra* based on CHELSA climatologies. Left, current; Right, future (2070). Future projections include two emission scenarios (SSP126 and SSP585) and three GCMs.

### Future climate suitability

At known populations (i.e. presence points), climate suitability will decline for both species according to both climate data sources, and for all GCMs and scenarios. The predicted decline is generally more severe when based on Worldclim data as compared to CHELSA (Fig. 5).

**Fig. 5.**
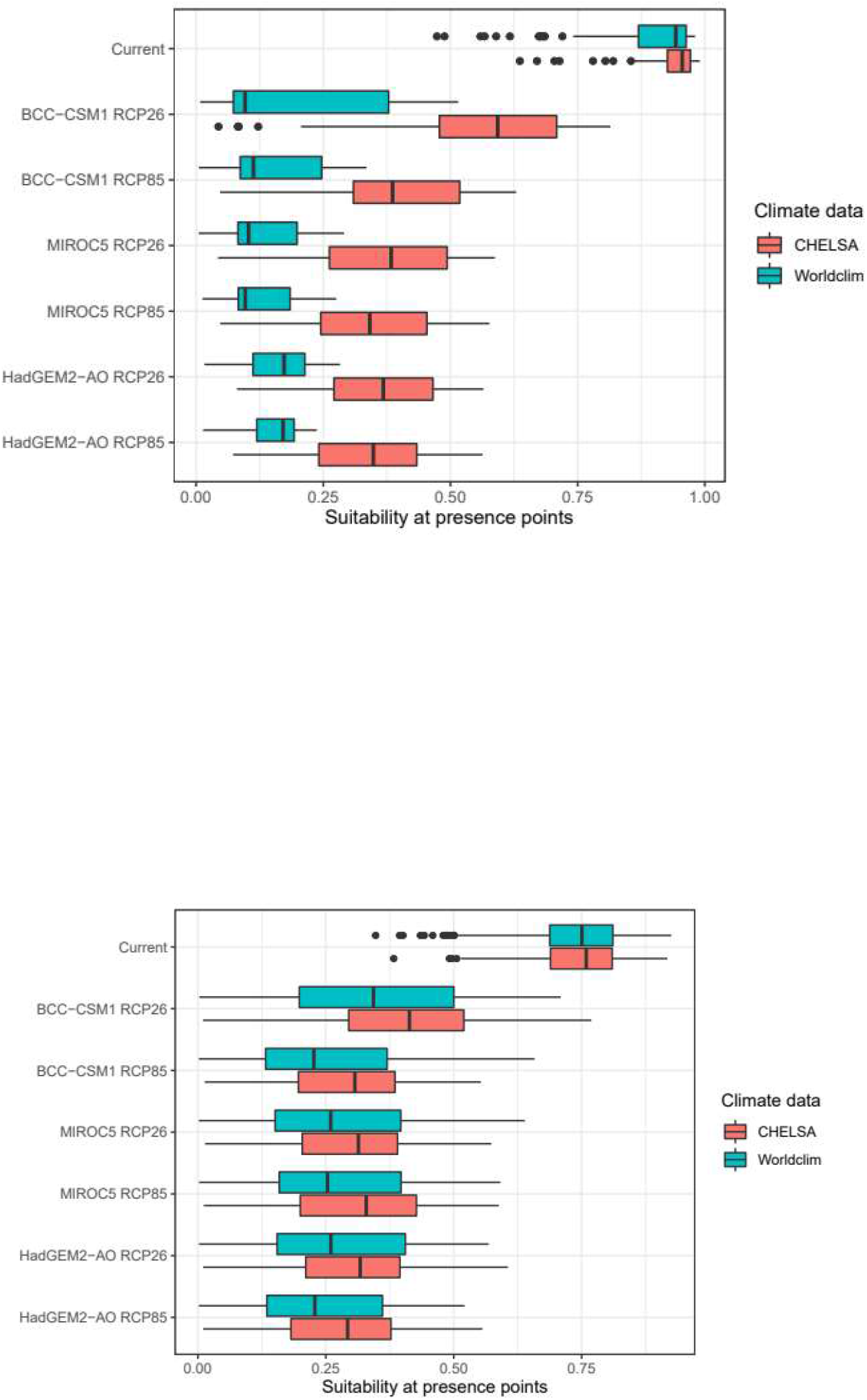
Current and future (2070) environmental suitability for two alpine salamanders (Top: *Salamandra lanzai*; Bottom: *S. atra*) according to two climate data sources (Worldclim and CHELSA) and a range of scenarios and GCMs.

For *S. lanzai*, models based on Worldclim predicted that the most suitable areas will be located in the eastern Alps. According to CHELSA, the most suitable areas will be located at the current distribution and in the northern part of central Alps. For *S. atra*, the most suitable areas will remain similar to the current ones, but with suboptimal conditions throughout the Alps.

### Accounting for uncertainty and competition

When taking into account the uncertainty related to model replicates and climate data (source, GCMs and scenarios), the most suitable areas for *S. lanzai* will be located in central Alps / eastern Switzerland, between the Swiss National Park and the Park Ela (Fig. 6). For *S. atra*, the most suitable areas also will be in central Alps, although further west to *S. lanzai*’s projected suitable area (i.e. around the Beverin Nature Park).

**Fig. 6.**
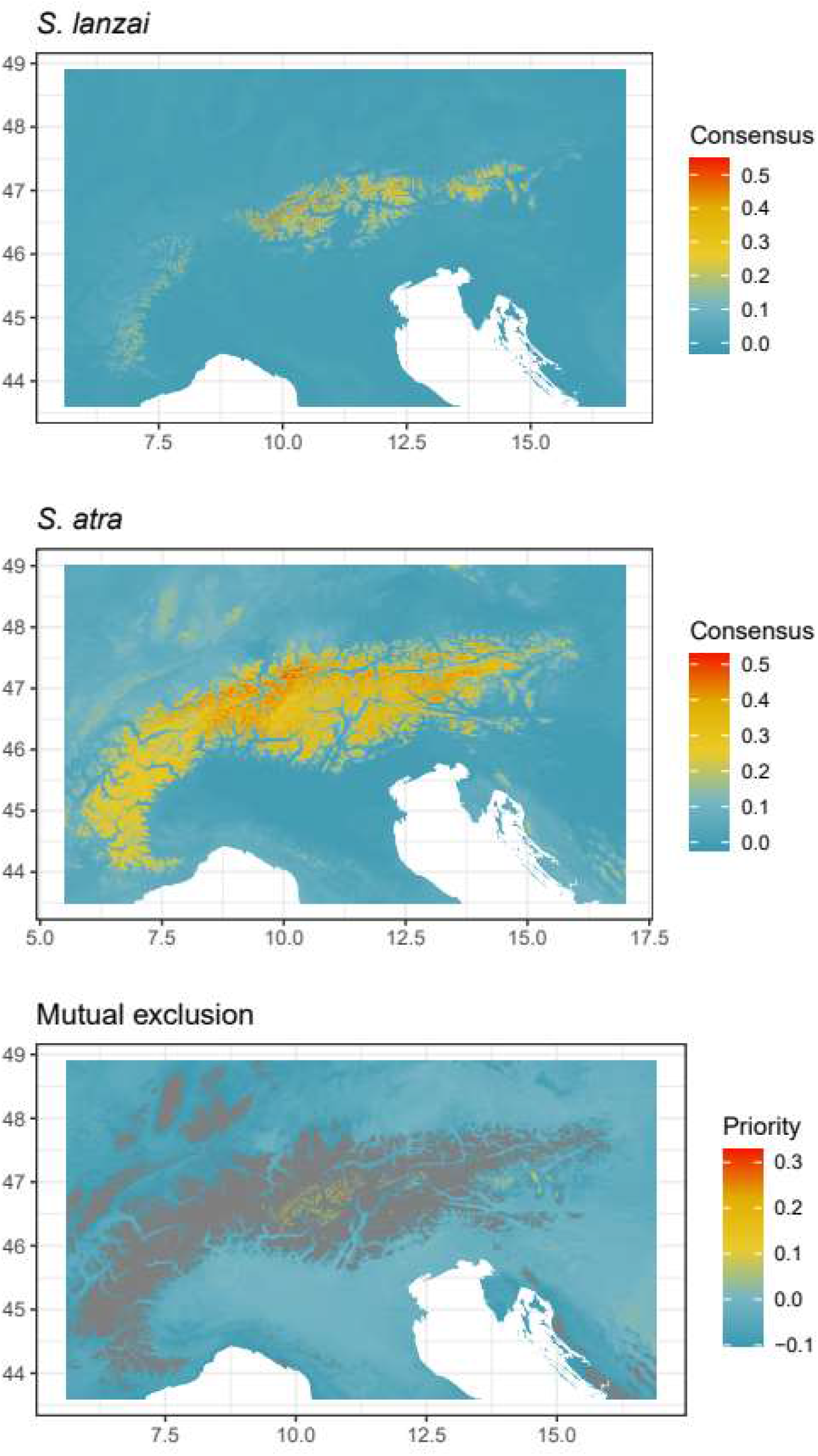
Consensus predictions for current climate suitability of *Salamandra lanzai* and *S. atra* (mean prediction across all models penalised by uncertainty), with the most suitable areas for *S. lanzai* excluding suitable areas for *S. atra*.

When accounting for potential competition with *S. atra*, the most suitable areas by 2070 will be located in the Italian Alps, in the Parco Nazionale dello Stelvio and the Parco delle Orobie Bergamasche. Our projections did not include the area currently occupied by *S. lanzai* because it is also projected to be suitable to *S. atra*. However, it is unlikely that *S. atra* would reach this region through natural dispersion. As such, the current distribution of *S. lanzai* may still be included in the priority areas for the conservation of this species.

## Discussion

We projected the future climate suitability of two Alpine salamanders to inform possible conservation guidelines which takes into account climate projection uncertainty and competition. Our results suggest a severe effect of climate change on both species. We provide a single conservation priority map as a practical tool for practitioners.

### Ecological niche models

*Salamandra lanzai* is found in cold areas. We were not able to identify a minimum temperature for activity, presumably because our presence data encompassed the coldest pixels of the selected background. Suitable areas are characterized by low temperature variation throughout the year and the day. Hot or short winters may increase metabolic costs and affect adult survival (Kissel et al., 2019), while extremely cold nights may affect larval and adult survival (Madison, 1997). Paradoxically, hot winters may expose salamanders to extreme cold because of lower snow cover in winter. Indeed, salamanders often use mammal shelters (including marmot and other rodents burrows; Ribéron and Miaud 1999) and consequently may suffer from thinner snow cover (due to less effective thermal isolation) as temperature increases (Tafani et al. 2013). According to Worldclim projections, *S. lanzai* occupies the wettest areas of the western Alps. In this case, winter precipitation explained better than summer precipitation the current distribution. This effect may constitute a proxy for the available amount of water stored in the soil for the rest of the year. Surprisingly, projections based on CHELSA identified areas that are among the driest of the study area. CHELSA takes into account orographic factors and wind exposure. *Salamandra lanzai* usually occupies microhabitats with little wind exposure, presumably because wind may exacerbate the effect of cold on salamanders, as well as increase the chances of desiccation through evapotranspiration. We interpret the effect of summer precipitation for CHELSA as a proxy for wind exposure, which may explain the shape of the relationship.

Our data also enabled us to identify an optimal temperature window for *S. atra* (1–7°C annual mean) which is consistent with our expectations regarding activity. The wetter the habitat during summer, the more likely it is to be occupied by the species. Results are less consistent regarding winter precipitation, with a notable disagreement between Worldclim and CHELSA projections. Differences may be related to the different treatment of topography between both climatologies. We call for further investigation on the interpretation of precipitation predictors in the Alps.

### On the use of 1 km-resolution climate data in mountains

The use of downscaled climate data is usually recommended in mountainous environments (Lembrechts et al., 2019). Recent studies of mountain-dwelling salamanders, including *S. atra*, used downscaled climatic predictors to better take into account the heterogeneity of mountain environments (Feldmeier et al., 2020; Stickley & Fraterrigo, 2023). Both studies found that accounting for micro-climate reduces the predicted impact of climate change on salamanders, notably due to buffering effects of forest vegetation and micro-climate availability in valleys non-exposed to the sun. In our case, both species live in forested areas (apart from the highest altitude populations of *S. lanzai* living in saxicolous habitats), suggesting that forest management might not be a relevant avenue for the conservation of alpine salamanders. The approach of Feldmeier et al. (2020) highlights the consideration of habitat management on north-exposed valleys for *S. atra*. Our results based CHELSA data—which take into account valley exposure (Karger et al., 2017)— enable us to examine the orientation of suitable valleys at 1 km-resolution, and fully support their conclusions since most of the future suitable areas are predicted in north-exposed valleys (Fig. 4).

Nonetheless, we stress that our future projections suggest a strong decline in climate suitability for *S. atra* also in these areas (Fig. 5). Very high resolution climate data are rarely available at large scale, and future projections often implies statistical downscaling based on transposed and interpolated anomaly between coarse and fine scale data (e.g., Feldmeier et al., 2020; Lembrechts et al., 2019). Despite the use of 1 km resolution omits the presence of microhabitats, our future projections are still meaningful when considered relative to predictions of current suitability.

### The future of Alpine salamanders

We predict an important decline in climate suitability for both species, with only suboptimal climates left available by 2070. Winter temperature rising may increase metabolic costs, which would affect survival rates. Warmer summers likely reduce activity for *S. atra* (Geiger 2006), which supports a lower foraging efficiency, reproduction and overall survival. Precipitations are projected to decrease, which may delay age at maturity, reduce body length and growth rate, and increase gestation duration (Miaud et al. 2001; Andreone et al. 2004). According to the source of climate data, *S. lanzai* may either persist within its current distribution area or go fully extinct. We recommend to maintain or improve conservation efforts within the current range, and consider translocation to the identified areas of central Alps as a possible avenue to be considered in the future for the most threatened populations. The results and efficiency of translocation action is variable (Seddon, Strauss, & Innes, 2012; Thévenin, 2019), and since both *S. lanzai* and *S. atra* are aplacental viviparous species, these should be carefully evaluated and their cost and benefits assessed. While, viviparity would assure a better isolation of foetuses within mother’s body, it is evident that the evolution of this K-oriented breeding strategy require habitat stability (i.e. caused either by climate warming or by translocation). In parallel to these evaluations we encourage to consider developing captive breeding and gametes cryopreservation as possible actions for conservation.

### Concluding remarks

This study showed the importance of considering multiple climatologies when projecting climate change effects on species distributions in a mountainous environment. We provide a high-resolution map accounting for uncertainty related to climate data, habitat requirements and biotic interactions (competition), which is here made available to practitioners to support the decisions-making process for the conservation actions targeting a Critically Endangered species potentially on the brink of extinction. We encourage the documentation of species with similar ecological requirements to improve the consideration of competition in species distribution models.

## Acknowledgements

We thank everyone that contributed to the data collection in the field. Portuguese National Funds through FCT (Fundação para a Ciência e a Tecnologia) support the research contract (2020.00823.CEECIND/CP1601/CT0003) and research project (PTDC/BIA-EVL/31254/2017) to Angelica Crottini and the research contract to Nicolas Dubos [ICETA_2021_26]. We are grateful to Lucio Bonato, Gentile Francesco Ficetola, Claude Miaud and Benedikt R. Schmidt for useful discussions. We thank Matteo Quartesan for data provision and useful exchanges.

## Author contributions

ND led the writing of the manuscript, designed and conducted the analyses. AH contributed to data analysis and writing. DS and FA have made substantial contribution to acquisition of the data. AC and FA have made a substantial contribution to data interpretation, conception of the study and manuscript revision. All authors have approved the final version for submission, have agreed to be accountable for the accuracy and integrity of the work that they conducted, have agreed to resolve queries relating to the work that they conducted and contributed to the study.

## Supporting information

**Fig. S1.**
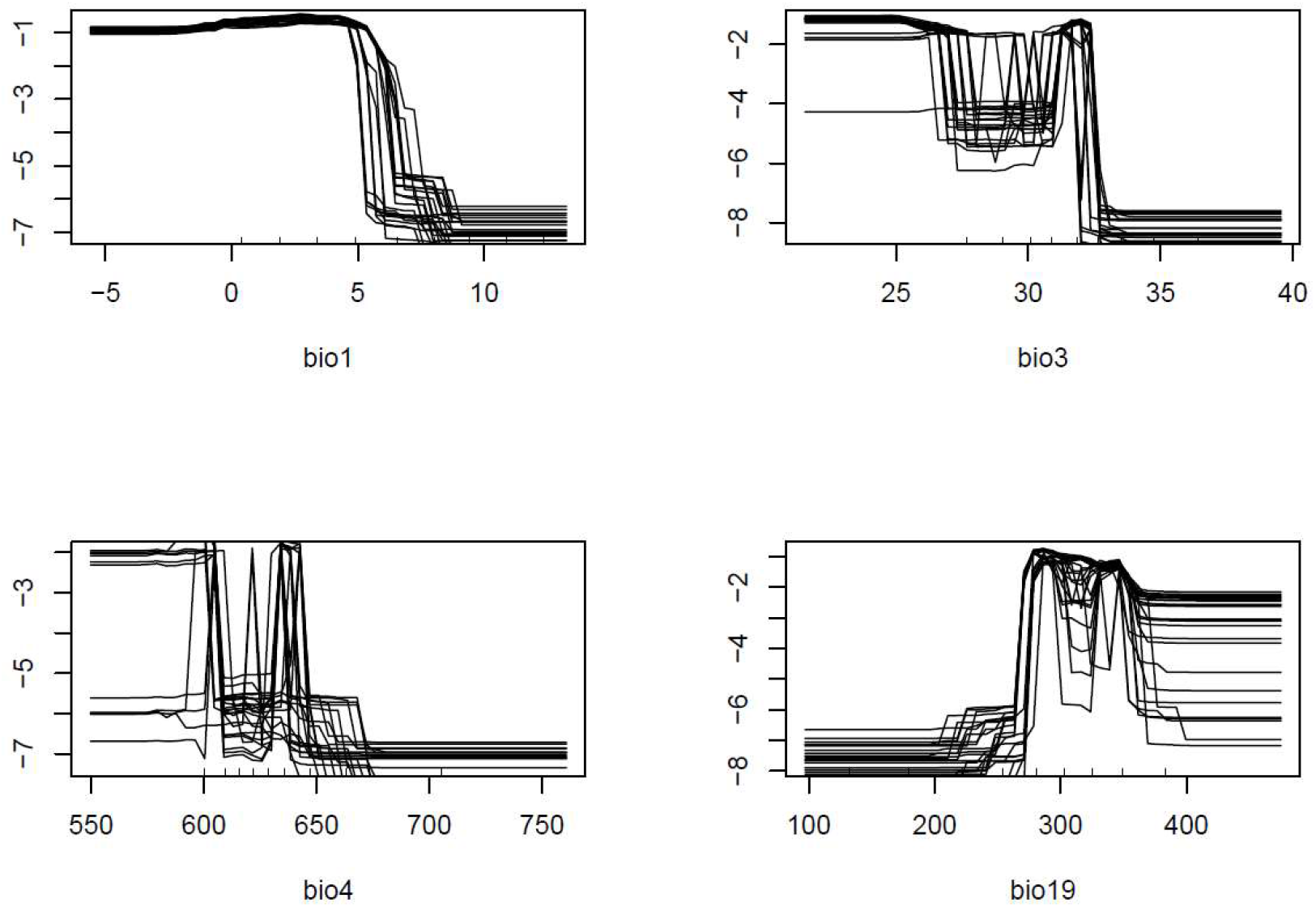
Predicted values for the climate suitability of *S. lanzai* based on Worldclim predictors

**Fig. S2.**
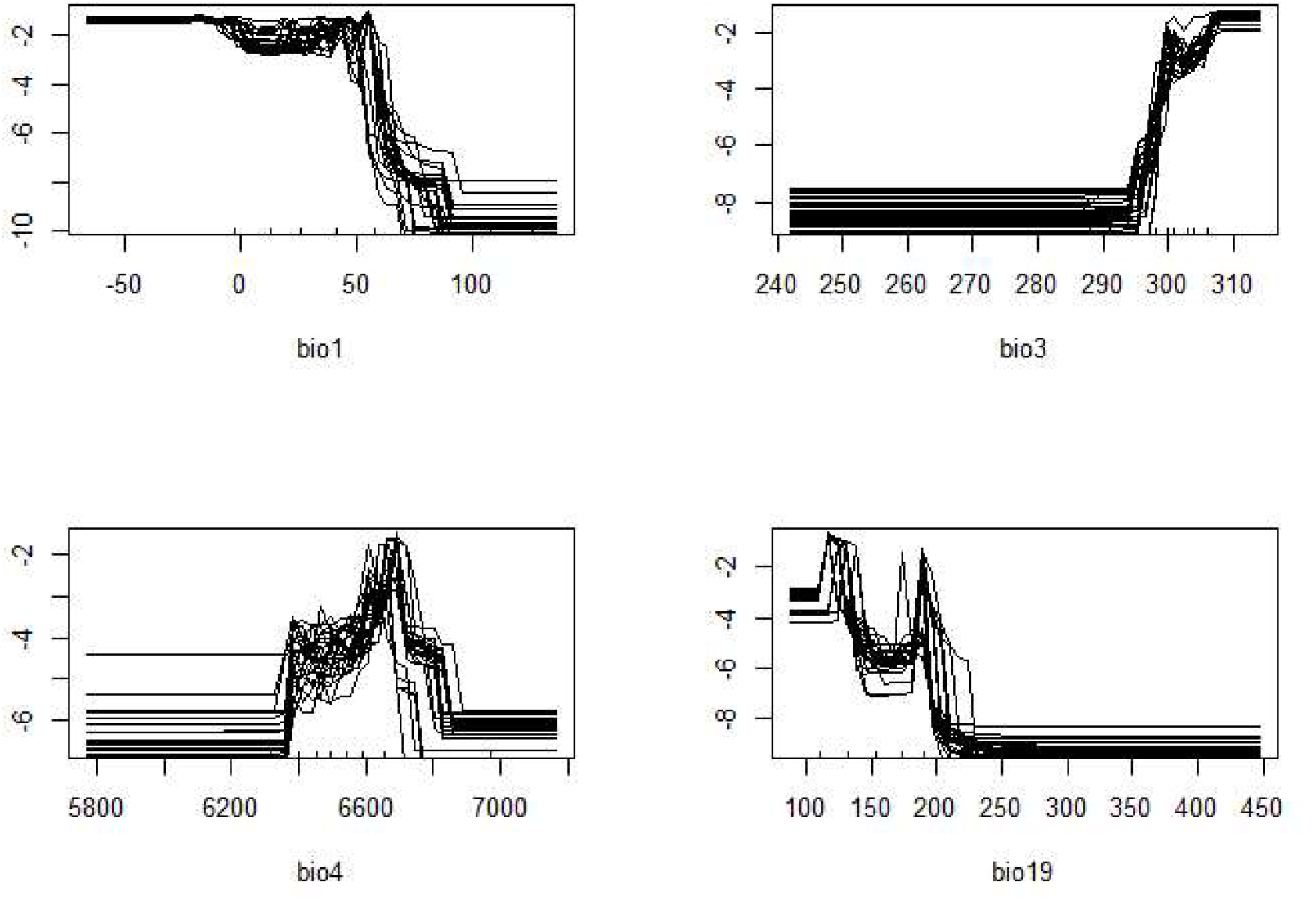
Predicted values for the climate suitability of *S. lanzai* based on CHELSA predictors

**Fig. S3.**
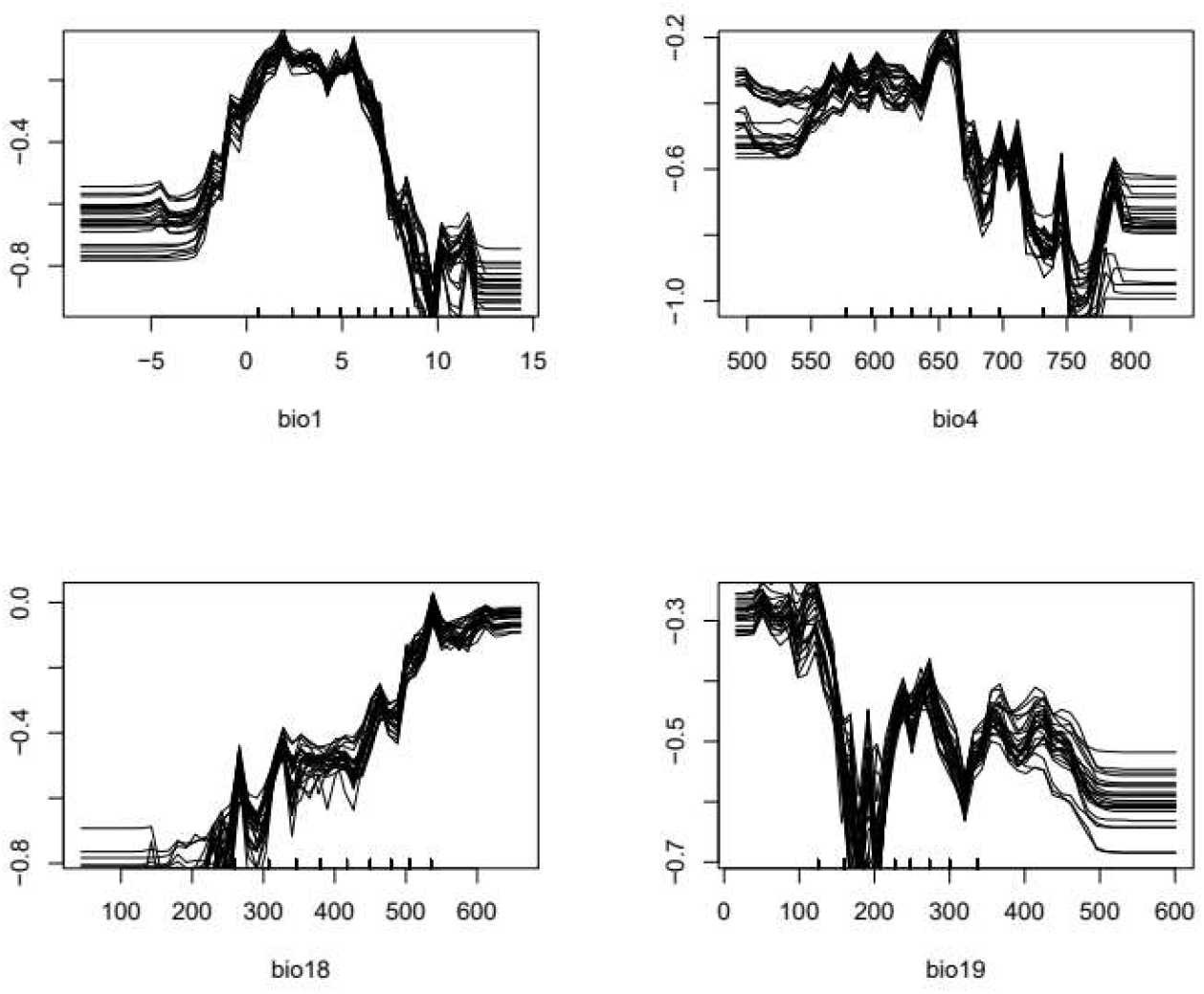
Predicted values for the climate suitability of *S. atra* based on Worldclim predictors

**Fig. S4.**
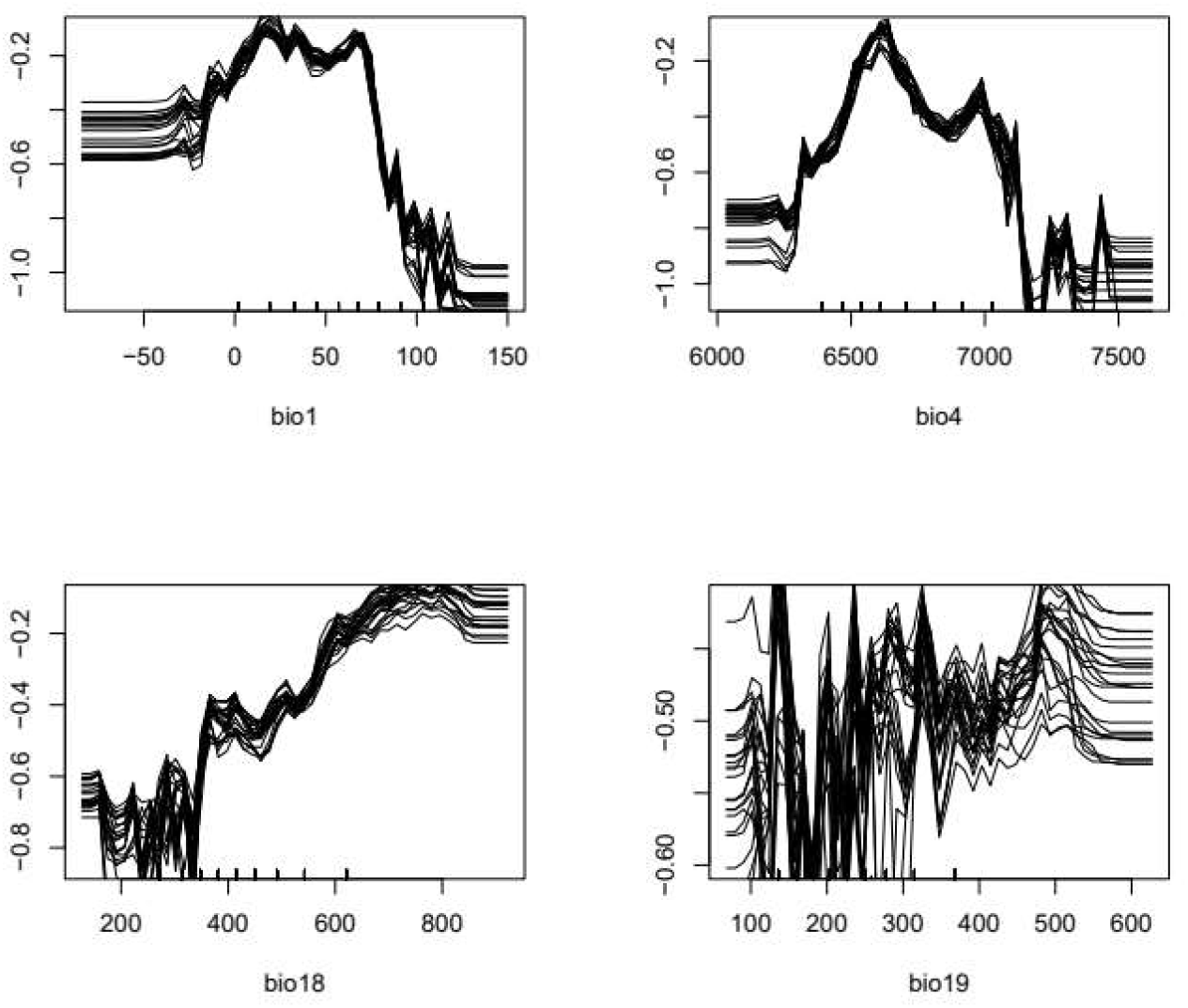
Predicted values for the climate suitability of *S. atra* based on CHELSA predictors

